# Intrinsic and TDP-43 dysfunction-induced catabolic stress elicit neuroprotective cellular degradation in ALS-vulnerable motor neurons

**DOI:** 10.1101/2025.05.06.652552

**Authors:** Kazuhide Asakawa, Takuya Tomita, Shinobu Shioya, Hiroshi Handa, Yasushi Saeki, Koichi Kawakami

## Abstract

Selective neuronal vulnerability is a defining feature of neurodegenerative disorders, exemplified by motor neuron degeneration in amyotrophic lateral sclerosis (ALS). The nature of motor neurons underlying this selectivity remains unresolved. Here, by monitoring autophagy at single-cell resolution across the translucent zebrafish spinal cord, we identify motor neurons as the cell population with the highest autophagic flux. Large spinal motor neurons (SMNs), most susceptible to ALS, exhibit higher flux compared to smaller SMNs and ALS-resistant ocular motor neurons. Notably, large SMNs accelerates both autophagy and proteasome-mediated degradation, which are further augmented by TDP-43 loss. Additionally, acceleration of multiple unfolded protein response pathways indicates their innate tendency to accumulate misfolded proteins. Enhanced cellular degradation in large SMNs is neuroprotective as its inhibition halts axon outgrowth. These findings propose that cell size-associated degradation load underlies selective neuronal vulnerability in ALS, highlighting the alleviation of catabolic stress as a target of therapy and prevention.

## Introduction

Cells constituting the central nervous system (CNS) exhibit varying degrees of susceptibility to various diseases. Amyotrophic lateral sclerosis (ALS), a currently incurable neurodegenerative disorder, exemplifies disease-specific neuronal susceptibility in which motor neurons responsible for muscle contraction undergo selective degeneration. Remarkably, selective vulnerability exists, even within this functionally similar neuronal population. For instance, the motor neurons innervating the extraocular muscles are resistant to ALS, though the underlying mechanisms remain poorly understood ^1–3^. Consequently, eye movements tend to be relatively preserved, even in the end-stage of the disease in ALS, whereas most bodily movements become progressively impaired. By contrast, large motor neurons are particularly vulnerable to ALS and normal aging ^4^ ^5, 6^. Large motor neurons, or α-motor neurons, are characterized by large cell bodies with highly branched dendritic arbors that receive a multitude of synaptic inputs. These neurons have a high conduction velocity due to their long axons with a large diameter and thick myelination and control large numbers of muscle fibers to generate strong voluntary body movements. However, the physiological and molecular mechanisms by which these cellular traits contribute to the selective vulnerability to ALS and aging remain unclear.

The accumulation of misfolded proteins or protein aggregates is a characteristic of most neurodegenerative diseases. Deposition of ubiquitin-positive inclusion bodies containing aggregates of the trans-activation response element (TAR) DNA-binding protein 43 (TDP-43) in motor neurons is a major hallmark of ALS ^7, 8^, suggesting that a decline in cellular degradation could be intrinsically linked to the as-yet-unidentified root causes of ALS. Macroautophagy (hereafter referred to as autophagy) is a critical cellular degradation mechanism for eliminating and recycling damaged proteins and organelles. Dysfunction of this process may impair neural function and result in neurodegeneration ^9, 10^. Human genetics have identified multiple autophagy-related genes as causative factors of neurodegenerative diseases, including ALS ^11–17^. The accumulation of autophagosomes, a double-membraned vesicles that engulf degradation substrates and deliver them to lysosomes, in spinal motor neurons (SMNs) has been reported in sporadic cases with ALS and animal models of the disease ^18–20^. These observations substantiate that autophagy-related cellular degradation plays a protective role in physiology and pathophysiology of motor neurons ^21, 22^, although autophagy can have adverse effects during certain stages of the disease course ^21^. In neuronal cells, autophagy operates constitutively at certain levels and the physiological role of such basal autophagic flux may differ from that of the canonical response to starvation ^23^. The involvement of autophagy in the regulation of motor neuron sub-compartments, including axon formation and synaptogenesis, has been reported in multiple models ^21, 24–27^. In addition to autophagy, the ubiquitin-proteasome system (UPS) is a key cellular degradation system involved in ALS pathogenesis. In mice, motor neuron-specific knockout of a proteasome subunit results in ALS phenotypes, whereas knockout of the autophagy-related gene ATG7 does not, suggesting that the UPS and autophagy may play distinct roles in ALS pathogenesis ^28^ ^29^. The UPS exhibits varying levels of activity in mouse embryonic stem cell-derived cranial motor neurons and SMNs, which correspond to ALS-resistant and ALS-vulnerable neurons, respectively ^3^, indicating a potential role for UPS activity in determining neuronal susceptibility to ALS. Currently, it remains unclear whether the activity of major cellular degradation pathways, such as autophagy and the UPS, differs across various CNS cell types and how such variations contribute to the differing neuronal vulnerability observed in ALS. To address these questions, comparing the dynamic aspects of cellular catabolism in vivo between ALS-vulnerable and ALS-resistant neurons is imperative. However, such real-time in vivo comparisons are not readily feasible, even in model animals, let alone in humans.

In the present study, we performed a cellular-resolution scanning of the autophagic flux across the translucent zebrafish spinal cord and identified motor neurons as the cell population with the highest level of autophagic flux. A comparison among the motor neuron subtypes revealed that large SMNs displayed higher autophagic flux than smaller SMNs as well as brainstem ocular motor neurons, both of which are relatively resistant to ALS. Remarkably, large SMNs not only exhibited enhanced autophagy but also demonstrated proteasome-mediated degradation. These processes were further intensified by the loss of the ALS-related protein TDP-43. The naturally activated multiple unfolded protein response pathways in large SMNs suggest an inherent tendency to accumulate misfolded proteins. These results posit that the cellular degradation load linked to cell size underlies motor neuron vulnerability and is a crucial determinant of selective susceptibility to ALS.

## Results

### Diversity of autophagy activity among neuronal cells in the spinal cord

Differential basal autophagic activity across various tissues has been revealed in zebrafish using the ratiometric autophagic flux probe GFP-LC3-RFP-LC3ΔG^30^. The probe assesses the stability of the key autophagosome marker, microtubule-associated protein light chain 3 (LC3) by releasing degradable GFP-LC3 and a non-degradable internal control, RFP-LC3ΔG through cleavage by the ATG4 protease upon autophagy activation and reflects autophagy flux acceleration by a decrease in the GFP-LC3/RFP-LC3ΔG ratio. Using this probe, the basal autophagic flux was demonstrated to be relatively lower in the spinal cord than in the skeletal muscle ^30^. To investigate the diversity of autophagic flux within the spinal cord at single-cell resolution, we generated transgenic zebrafish (Tg[UAS:GFP-LC3-RFP-LC3ΔG]) in which GFP-LC3-RFP-LC3ΔG can be expressed ubiquitously or in a defined cell type in a Gal4-dependent manner (Fig. 1a). First, we broadly expressed GFP-LC3-RFP-LC3ΔG in the spinal cord using the ubiquitous Gal4 driver Tg[SAGFF73A] ^31^. Expression levels of GFP-LC3 and RFP-LC3ΔG varied among the spinal cord cells in Tg[SAGFF73A] Tg[UAS:GFP-LC3-RFP-LC3ΔG] double transgenic fish (Fig. 1b, Supplementary Video 1). We noticed a cell population with oval-shaped cell bodies and a markedly low GFP-LC3/RFP-LC3ΔG ratio, indicating high autophagic flux, in the ventrolateral side of the spinal cord at 30 hours post-fertilization (hpf) (Fig. 1b). Neuronal populations in the spinal cord commonly exhibit spatially characteristic clustering according to cell type ^32^. Neurons responsible for sensory functions are primarily positioned dorsally, whereas those controlling motor execution are predominantly located ventrally. Thus, we hypothesized that the ventral cells with a low GFP-LC3/RFP-LC3ΔG ratio include SMNs, which cluster to form a motor column on the ventral side of the spinal cord in zebrafish ^33, 34^. To address this possibility, we restricted GFP-LC3-RFP-LC3ΔG expression to the caudal primary motor neurons (CaPs) in the ventral side and tactile sensing Rohon-Beard (RB) cells in the dorsal side of the spinal cord, using the Gal4 driver Tg[SAIG213A] ^35^ (Fig. 1c). We found that GFP-LC3/RFP-LC3ΔG values in the soma regions were significantly lower in the CaPs than in RB cells at 30 hpf (Fig. 1d), showing that CaPs have a higher level of basal autophagic flux than the RB cells. The Tg[SAIG213A] driver also labeled some types of interneurons with diverse cell morphologies. The GFP-LC3/RFP-LC3ΔG ratio in these interneurons was intermediate between those of observed in CaPs and RB cells (Fig. 1d). Taken together, these observations suggest that the cell populations with high autophagic flux in the ventrolateral spinal cord includes SMNs. Zebrafish assembles a neural circuit for touch-evoked swimming movements as early as by 21 hpf, where tactile afferent signals elicited from RB cells are transmitted to SMNs, including CaPs ^36, 37^. Therefore, these results demonstrate that the functional neural circuit for sensory-evoked locomotion consists of cells with different levels of autophagic flux.

**Fig. 1.**
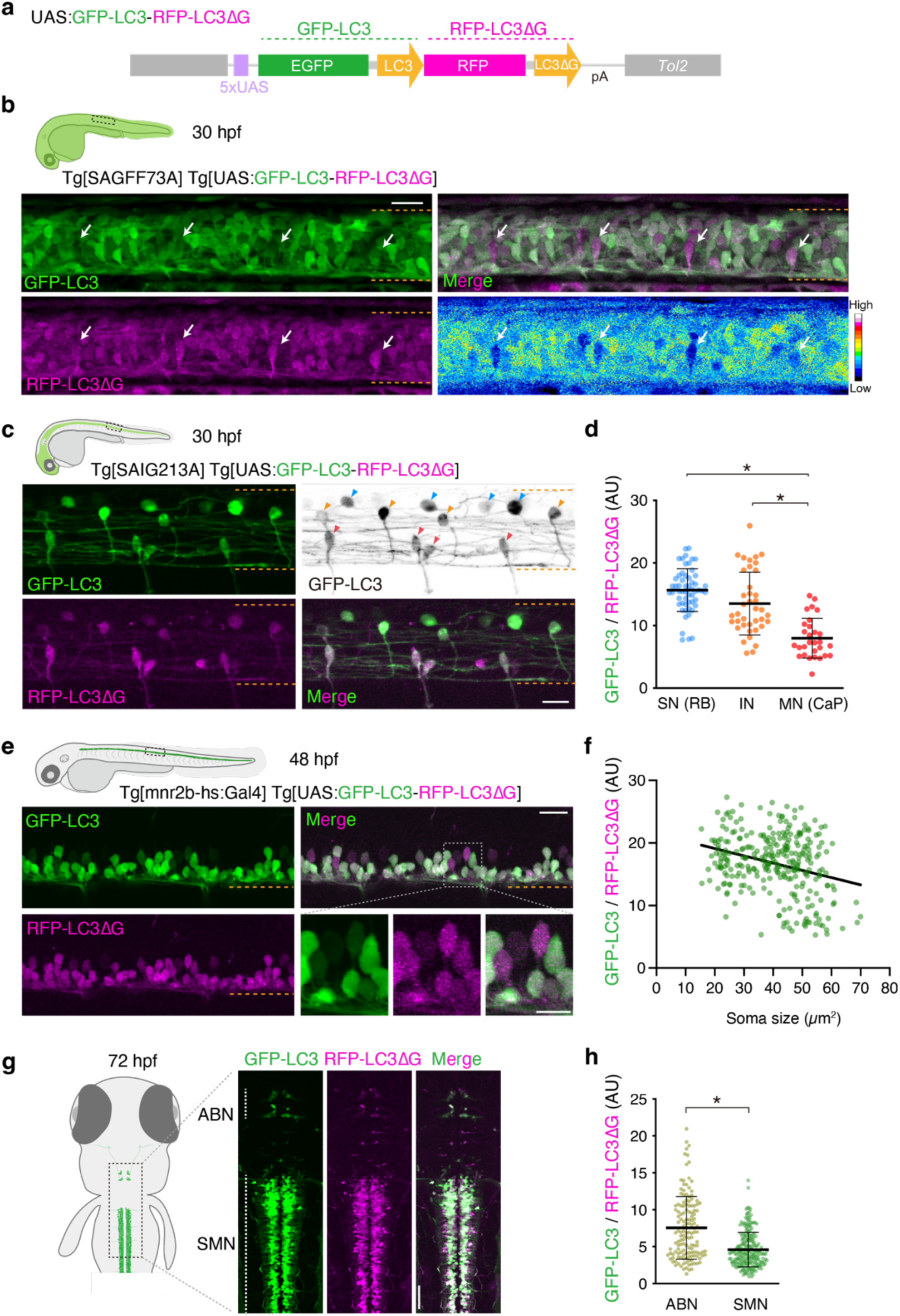
Autophagic flux scanning across spinal cord neurons. (a) Structure of Tg[UAS:GFP-LC3-RFP-LC3ΔG]. (b) Lateral view of the spinal cord of Tg[SAGFF73A] Tg[UAS:GFP-LC3-RFP-LC3ΔG] fish at 30 hpf. The arrows indicate the representative cells displaying low GFP-LC3/RFP-LC3ΔG ratios. (c) Lateral view of the spinal cord of Tg[SAIG213A] Tg[UAS:GFP-LC3-RFP-LC3ΔG] fish at 30 hpf. The arrowheads indicate sensory neurons (blue, RB cells), interneurons (orange), and motor neurons (red). (d) GFP-LC3/RFP-LC3ΔG ratios in sensory neurons (SN, RB cells, N=56), interneurons (IN, N=29), and motor neurons (MN,CaP, N=39). Data were obtained from four animals. *, p < 0.0001 (unpaired t-test, two-tailed). (e) Lateral view of the spinal cord of Tg[mnr2b-hs:Gal4] Tg[UAS:GFP-LC3-RFP-LC3ΔG] fish at 48 hpf. (f) Dot plots of GFP-LC3/RFP-LC3ΔG ratios as a function of soma size (cross-sectional area). N=293 *mnr2b*+ cells from 12 animals. (g) Dorsal view of the brainstem‒spinal cord junction of Tg[mnr2b-hs:Gal4] Tg[UAS:GFP-LC3-RFP-LC3ΔG] fish at 72 hpf. (h) Dot plots of GFP-LC3/RFP-LC3ΔG ratios of ABNs (N=142 cells) and SMNs (N=244 cells). Data were obtained from seven animals. Dashed orange lines indicate the dorsal and ventral limit of the spinal cord. *p < 0.0001 (unpaired two-tailed t-test). Error bars show SD. The bars indicate 20 µm (B, C, larger panels in E), 10 µm (smaller panels in E), and 50 µm (G).

### Inverse correlation between basal autophagic flux and cell size of SMNs

Having discovered that SMNs exhibit the highest level of autophagic flux in the spinal cord, we further explored whether the autophagic flux varies among motor neuron subtypes. The SMNs represent a heterogeneous population with diverse cell sizes corresponding to their distinct physiological roles and connectivity patterns ^38^. The fact that CaPs represent one of the largest SMN types in zebrafish ^39^ prompted us to examine the relationship between cell size and autophagic flux intensity in SMNs. Using the pan-motor neuronal Gal4 driver Tg[mnr2b-hs:Gal4] ^40^, we expressed GFP-LC3-RFP-LC3ΔG in most of the SMNs and measured the GFP-LC3/RFP-LC3ΔG ratio in their soma regions at 48–50 hpf (Fig. 1e). Intriguingly, by plotting the GFP-LC3/RFP-LC3ΔG values against soma size, we observed a significant trend that the GFP-LC3/RFP-LC3ΔG ratio decreased as soma size increased (Fig. 1f). This indicates that large SMNs tend to have more enhanced autophagic flux than small SMNs.

Motor neurons are the primary neuronal type that degenerates in the motor neuron disease ALS, with large SMNs being more vulnerable than smaller SMNs ^4^ ^5^. Enhanced autophagic flux in the large SMNs of zebrafish suggests a correlation between the intensity of autophagic flux and neuronal vulnerability to ALS. To further explore this idea, we compared the autophagic flux between SMNs and ocular motor neurons in the brainstem, which are known to be resistant to ALS. Purposing this, we took advantage of Gal4 expression in the abducens motor neurons (ABNs) of the brainstem in the Tg[mnr2b-hs:Gal4] line (Fig. 1g) ^40^. The GFP-LC3/RFP-LC3ΔG values of the ABNs and SMNs were compared in the same confocal sections including the junction between the brainstem and spinal cord at 72 hpf, when most ABNs had already connected to the lateral rectus muscle ^40^. We noted that the GFP-LC3/RFP-LC3ΔG values in the somas were highly variable in ABNs compared with those in SMNs (Fig. 1h). Nonetheless, ABNs displayed higher GFP-LC3/RFP-LC3ΔG values than SMNs, indicating that autophagic flux was generally lower in the ABNs than in SMNs. These observations further highlighted the correlation between enhanced autophagic flux and neuronal vulnerability in ALS.

### Loss of TDP-43 function accelerates basal autophagic flux in the SMNs

To explore the significance of accelerated autophagy in large SMNs in ALS, we examined how an ALS-related mutations affect autophagic flux in these neurons. We focused on the *tardbp/TARDBP* gene which encodes TDP-43 protein, mutations and protein aggregation of which are associated with most cases of ALS ^7, 8, 41^. Zebrafish have two genes that encodes TDP-43, *tardbp* and *tardbpl*. Using frame-shift mutations that introduce premature termination codons in both genes ^42^, we expressed the GFP-LC3-RFP-LC3ΔG reporter in the CaPs, RB cells, and interneurons of the TDP-43 double knockout (DKO) fish. Intriguingly, we found that the GFP-LC3/RFP-LC3ΔG values were significantly reduced in each of these neuronal types in the TDP-43 DKO fish at 48 hpf (Fig. 2a), showing that the loss of TDP-43 function accelerates autophagic flux regardless of neuronal type. This observation is consistent with the notion that TDP-43 dysfunction can lead to the production of abnormal proteins from mis-spliced RNA ^43^, imposing additional cellular degradation stress and accelerating the neuroprotective autophagic flux in SMNs. Furthermore, these results suggest that while TDP-43 dysfunction accelerates autophagic flux, the function of TDP-43 itself is not essential for this acceleration in large SMNs.

**Fig. 2.**
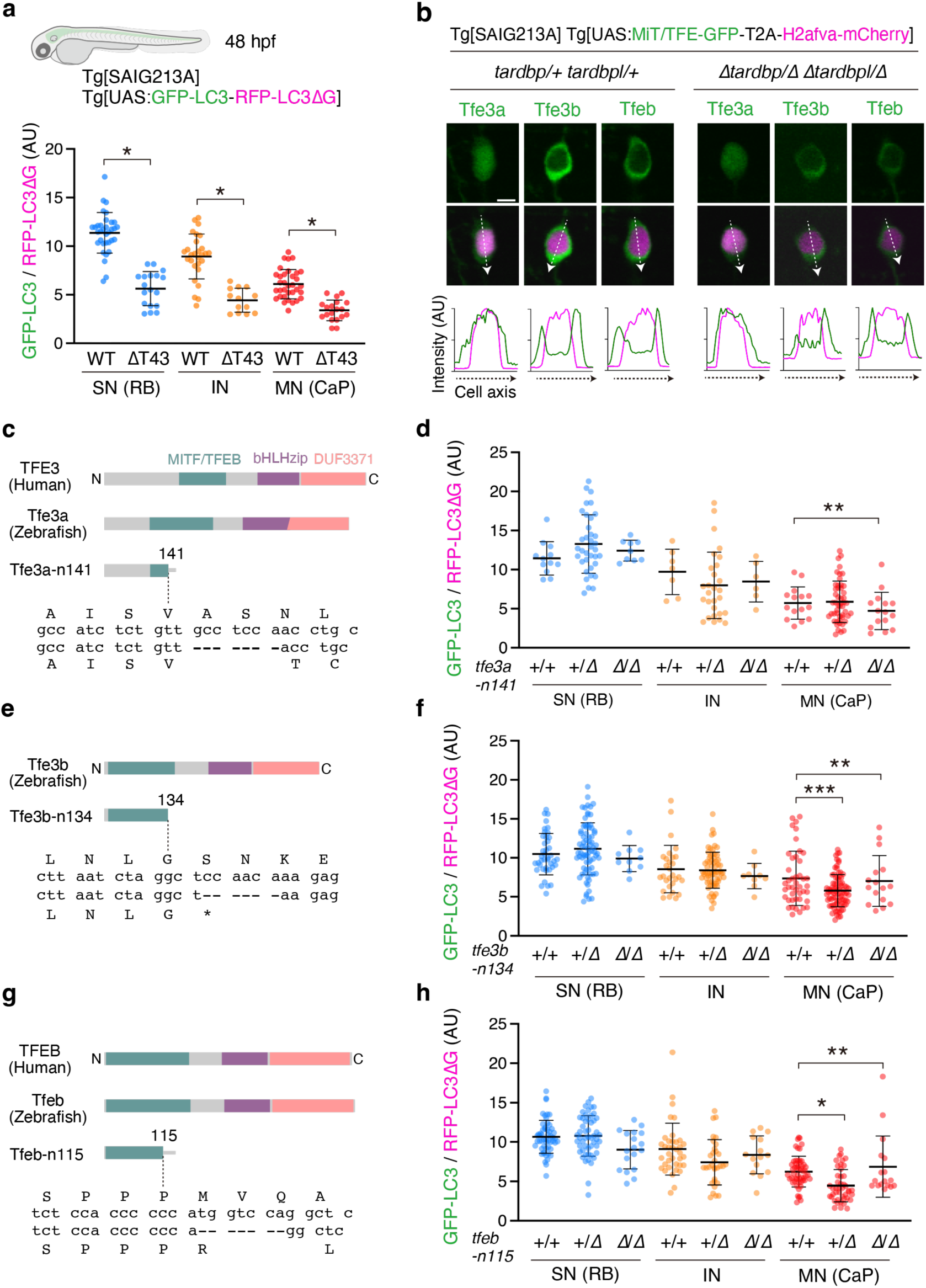
Loss of TDP-43 function accelerates autophagic flux in the SMNs. (a) GFP-LC3/RFP-LC3ΔG ratios in sensory neurons (SNs, RB cells), interneurons (INs), and motor neurons (MNs, CaPs) in wild-type (*tardbp*/*+ tardbpl*/*+*) and TDP-43 DKO (ΔT43, *tardbp-n115*/ *tardbp-n115 tardbpl-n94*/*tardbpl-n94*) backgrounds at 48 hpf. Data were obtained from eight WT fish (34 SNs, 28 INs, and 34 MNs) and four TDP-43 DKO fish (18 SNs, 13 INs, and 19 MNs). * p < 0.0001 (unpaired t-test, two-tailed). Error bars show SD. (b) Localization of GFP-tagged MiT/TFE in the CaPs of wild-type fish (left, *tardbp*/*+ tardbpl*/*+*) and TDP-43 DKO fish (*tardbp-n115*/ *tardbp-n115 tardbpl-n94*/*tardbpl-n94*) at 48 hpf. Fluorescence intensities of GFP and H2afva-mCherry along the cell axes (dashed arrows) are shown. The bar indicates 5 µm. (c) Structure of zebrafish Tfe3a protein and position of the *tfe3a-n141* mutation. MITF/TFEB: N-terminus conserved region, bHLHzip: basic Helix-Loop-Helix-zipper domain found in TFEB, DUF3371: Conserved domain of unknown function. (d) GFP-LC3/RFP-LC3ΔG ratios in three wild-type (*+/+*, 12 SNs, 7 INs, and 34 MNs), thirteen heterozygous (*+/Δ*, 36 SNs, 28 INs, and 50 MNs), and five homozygous (*Δ/Δ*, 9 SNs, 6 INs, and 15 MNs) *tfe3a-n141* mutant fish at 48 hpf. (e) Structure of zebrafish Tfe3b protein and position of the *tfe3b-n134* mutation. (f) GFP-LC3/RFP-LC3ΔG ratios in thirteen wild-type (*+/+*, 42 SNs, 29 INs, and 44 MNs), 22 heterozygous (*+/Δ*, 77 SNs, 68 INs, and 85 MNs), and four homozygous (*Δ/Δ*, 9 SNs, 9 INs, and 16 MNs) *tfe3b-n134* mutant fish at 48 hpf. (g) Structure of zebrafish Tfeb protein and position of the *tfeb-n115* mutation. (h) GFP-LC3/RFP-LC3ΔG ratios in 19 wild-type (*+/+*, 54 SNs, 36 INs, and 56 MNs), 25 heterozygous (*+/Δ*, 53 SNs, 33 INs, and 53 MNs), and 8 homozygous (*Δ/Δ*, 21 SNs, 17 INs, and 19 MNs) *tfeb-n115* mutant fish at 48 hpf. * p < 0.0001, ** p > 0.2, *** p = 0.0017 (unpaired t-test, two-tailed). Error bars show SD.

### Autophagic acceleration is independent of MiT/TFE transcription factors

Next, we investigated the potential upstream mechanisms driving the sustained, cell-type-specific activation of autophagy in large SMNs. In cultured human cells (HEK 293 and HeLa cells) ^44^, the knockdown of TDP-43 results in the nuclear accumulation of the MiT/TFE transcription factor TFEB ^44^, which enhances global gene expression in the autophagy–lysosome pathway (ALP) ^45, 46^. To determine the contribution of MiT/TFE transcription factors to autophagic flux acceleration in SMNs *in vivo*, we first examined the subcellular localization of the MiT/TFE transcription factors Tfeb, Tfe3a, and Tfe3b in CaPs. Each of these proteins was tagged with an enhanced green fluorescent protein (EGFP) and expressed along with the nuclear marker mCherry-tagged histone H2A (H2az2a-mCherry) using the Tg[SAIG213A] driver. Among the three transcription factors, only Tfe3a-EGFP was predominantly enriched in the nucleus, whereas Tfeb-EGFP and Tfe3b-EGFP were mainly localized in the cytoplasm at 48 hpf (Fig. 2b). The overall subcellular distribution patterns of these GFP-tagged proteins remained unchanged in TDP-43 DKO fish, suggesting that Tfe3a is a candidate transcription factor responsible for the acceleration of autophagic flux in SMNs. Next, to determine the causal relationships between the subcellular localization patterns of the MiT/TFE transcription factors and accelerated autophagy in SMNs, we introduced a frameshift mutation that resulted in premature termination in the MiT/TFE domain at the amino terminus of the *tfeb*, *tfe3a*, and *tfe3b* genes using the CRISPR-Cas9 system (*tfeb-n115, tfe3a-n141* and *tfe3b-n134*, respectively). However, contrary to our expectations, however, the homozygous *tfe3a-n141* mutation did not affect autophagic flux in CaPs or other neuronal cell types (Fig. 2c, d), indicating that nuclear enrichment of Tfe3a did not contribute to autophagy enhancement in CaPs. Consistent with the predominant cytoplasmic localization of Tfe3b-EGFP and Tfeb-EGFP, the homozygous *tfe3b-n134* mutation or *tfeb-n115* mutation did not cause a significant change in autophagic flux in CaPs (Fig. 2e–h). These observations collectively suggest that MiT/TFE transcription factors and the canonical ALP that they regulate play little to no role, if any, in accelerating autophagic flux in SMNs. We noted that heterozygous carriers of the *tfe3b-n134* or *tfeb-n115* mutations displayed a slight but significant acceleration of autophagic flux in CaPs, implying a dominant effect on the wild-type allele in this process. However, the mechanisms underlying these effects remain to be determined.

### The UPS is accelerated in large SMNs

To further explore why autophagic flux is enhanced in large SMNs, we investigated whether autophagy is particularly activated relative to other cellular degradation pathways. We specifically investigated the status of the ubiquitin-proteasome system (UPS), another major cellular degradation system. To this end, we developed a fluorescence-based reporter with a moderate half-life, EGFP-d410m, in which EGFP was tagged with a degron sequence (amino acids 410–461) from mouse ornithine decarboxylase carrying the T436A mutation, extending its half-life by 2-fold ^47^ (Fig. 4a, Extended Data Fig.1a). EGFP-d410m-T436A was stoichiometrically expressed alongside the internal control, mCherry, through self-cleaving P2A peptide-dependent translation. When expressed using the Tg[SAIG213A] driver, the EGFP-d410m-T436A/mCherry ratio significantly increased following bath treatment with the proteasome inhibitor MG132 (Extended Data Fig.1b), confirming that EGFP-d410m-T436A was indeed degraded via the UPS. We found that the EGFP-d410m-T436A/mCherry ratio was significantly lower in CaPs than in RB cells or interneurons (Fig. 3b, c), indicating that the UPS is intrinsically more active in SMNs than in other types of neuronal cells. Furthermore, among the SMNs, the EGFP-d410m-T436A/mCherry ratio significantly decreased as soma size increased (Fig. 3d, e), demonstrating that the UPS was significantly active in larger SMNs. Notably, under TDP-43 DKO conditions, the EGFP-d410m-T436A/mCherry ratio was significantly reduced in CaPs, as well as in RB cells and interneurons, indicating an acceleration of the UPS (Fig. 3c). We noted that TDP-43 disruption led to a reduction in the EGFP-d410m-T436A signal in most CaP cells. However, a subset of cells exhibited a striking increase, suggesting that the loss of TDP-43 function may have a bidirectional effect, markedly enhancing UPS activity in certain cells, while impairing it in others. Together, these results indicate that in SMNs, both autophagy and UPS are accelerated under normal and loss-of-TDP-43 function conditions, suggesting a general activation of intracellular degradation systems in large SMNs.

**Fig. 3.**
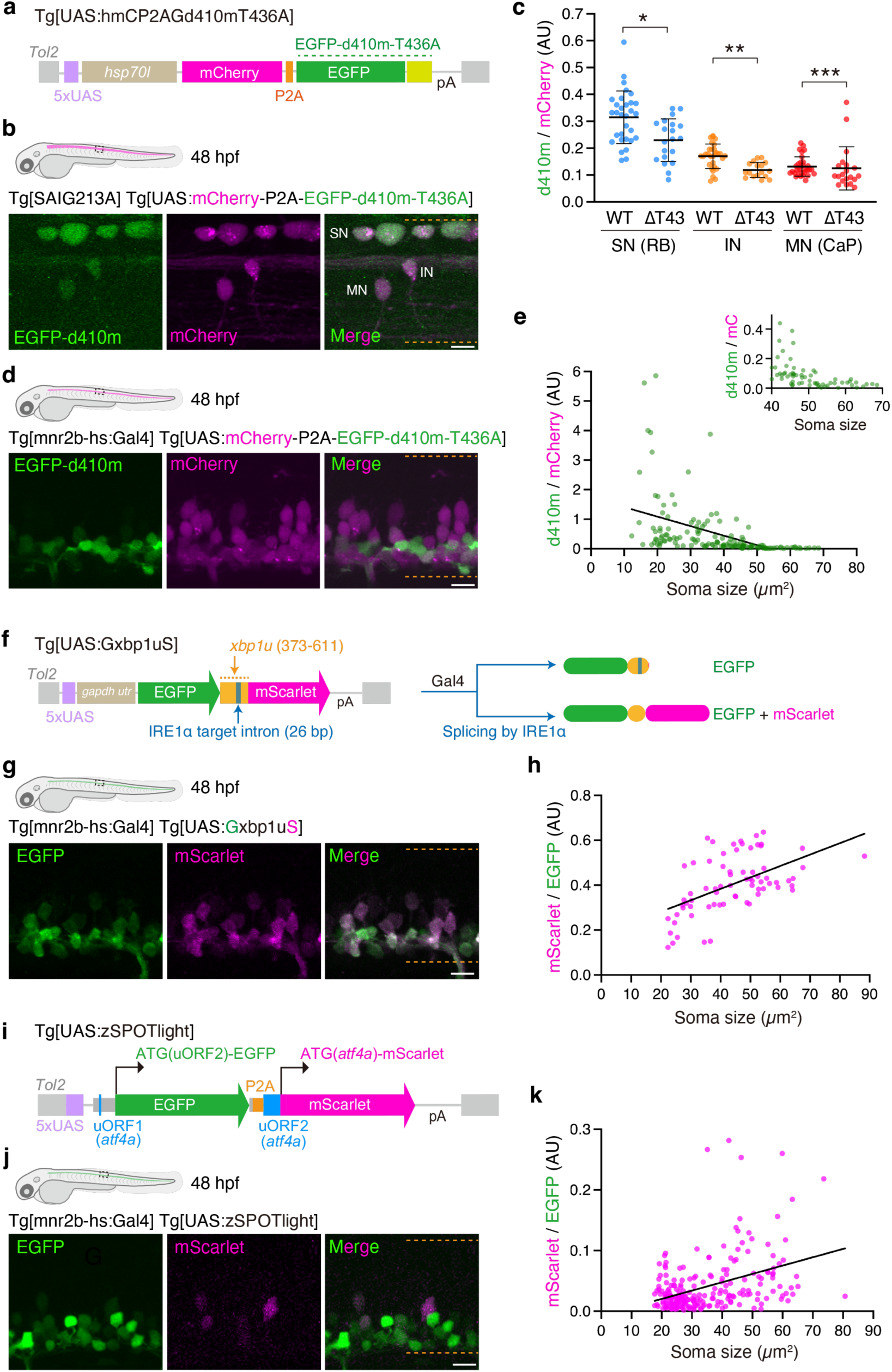
Enhanced UPS and UPR activities in large SMNs. (a) Structure of Tg[UAS:hmCP2AGd410mT436A]. (b) Lateral view of the spinal cord of Tg[SAIG213A] Tg[UAS:mCherry-P2A-EGFP-d410m-T436A] fish at 48 hpf. (c) GFP/mCherry ratios in sensory neurons (SNs, RB cells), interneurons (INs), and motor neurons (MNs, CaPs) in wild-type (*tardbp*/*+ tardbpl*/*+*) and TDP-43 DKO backgrounds at 48 hpf. Data were obtained from eight WT fish (31 SNs, 30 INs, and 31 MNs) and five TDP-43 DKO fish (20 SNs, 17 INs, and 20 MNs). * p = 0.002, ** p = 0.0001 (unpaired t-test, two-tailed), *** p = 0.0391 (Mann-Whitney test). Error bars show SD. (d) Lateral view of the spinal cord of Tg[mnr2b-hs:Gal4] Tg[UAS:mCherry-P2A-EGFP-d410m-T436A] fish at 48 hpf. (e) GFP/mCherry ratios in *mnr2b*-positive SMNs at 48 hpf. Data were obtained 146 cells from six fish. The slope was significantly non-zero (p < 0.0001). The magnification of GFP/mCherry was shown for larger SMNs (top, right). (f) Structure of Tg[UAS:Gxbp1uS]. An EGFP-mScarlet fusion protein is translated following Gal4-dependent transcription and IRE1α-dependent splicing. The 5’ untranslated region (500 bp) of the *gapdh* gene (*gapdh utr*) was used to enhance the Gal4-mediated expression. (g) Lateral view of the spinal cord of Tg[mnr2b-hs:Gal4] Tg[UAS:Gxbp1uS] fish at 48 hpf. (h) mScarlet/GFP ratios in *mnr2b*-positive SMNs at 48 hpf. Data were obtained from 72 cells across five fish. The slope was significantly non-zero (p < 0.0001). (i) Structure of Tg[UAS:zSPOTlight]. EGFP and mScarlet were translated from the initiation codon of uORF2 and *atf4a*, respectively. (j) Lateral view of the spinal cord of Tg[mnr2b-hs:Gal4] Tg[UAS:Gxbp1uS] fish at 48 hpf. (k) mScarlet/GFP ratios in *mnr2b*-positive SMNs at 48 hpf. Data were obtained 232 cells from five fish. The slope was significantly non-zero (p < 0.0001). Dashed orange lines indicate the dorsal and ventral limit of the spinal cord. Error bars show SD. The bars indicate 10 µm (B, D, G, J).

**Fig. 4.**
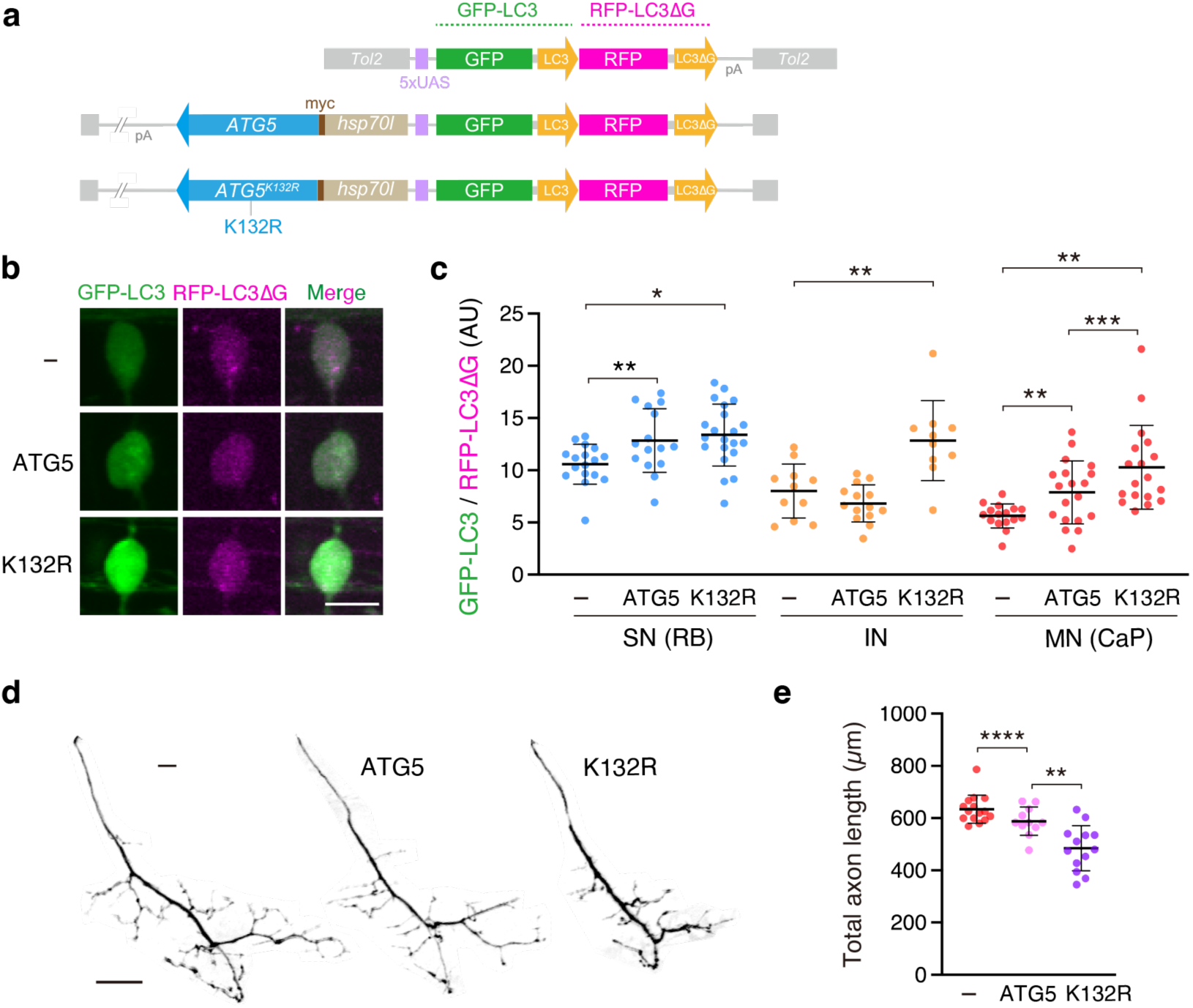
Autophagic acceleration is neuroprotective in large SMNs. (a) Structures of the bidirectional UAS transgenes for ATG5 (middle) and ATG5^K132R^ (bottom) expression with GFP-LC3-RFP-LC3ΔG. (b) Somas of the CaPs expressing GFP-LC3-RFP-LC3ΔG alone (-, top), the wild-type ATG5 (middle), and ATG5 ^K132R^ mutant (bottom). (c) GFP-LC3/RFP-LC3ΔG ratios in sensory neurons (SNs, RB cells), interneurons (INs), and motor neurons (MNs, CaPs). Data were obtained from three (wild-type, > 10 cells for each), five (ATG5, > 13 cells for each) and six (ATG5 ^K132R^, > 13 cells for each) animals. (d) Motor axons of CaPs expressing GFP-LC3-RFP-LC3ΔG alone (-, left), the wild-type ATG5 (middle), and ATG5 ^K132R^ mutant (right). (e) Total axon lengths of the CaPs in (D). Data were obtained from four (wild-type, 15 cells), three (ATG5, 11 cells) and four (ATG5 ^K132R^, 13 cells) animals. * p = 0.025, ** p < 0.001, *** p = 0.046, **** p = 0.0419, ***** p < 0.0001 (unpaired t-test, two-tailed). Error bars show SD. The bars indicate 10 µm (B), 20 µm (D).

### Intrinsic propensity of large SMNs to accumulate misfolded proteins

A possible cause for the inherently and generally elevated cellular degradation activity in large SMNs could be an increased accumulation of degradation substrates, such as misfolded proteins. To test this possibility, we investigated the status of the unfolded protein response (UPR), a cellular stress response mechanism that is activated when misfolded or unfolded proteins accumulate in the endoplasmic reticulum (ER). The activity of the three UPR sensors located in the ER membrane (IRE1α, PERK, and ATF6) is modulated by different specific factors ^48^. We first investigated the activity of the IRE1α-Xbp1 pathway by constructing a UAS reporter line (Tg[UAS:Gxbp1uS]). In this reporter, EGFP and mScarlet were fused via an ER stress-dependent *xbp1* intron of 26 base pairs (Fig. 3f). An EGFP-Xbp1-mScarlet fusion protein would be expressed when the stress-dependent *xbp1* intron was removed by IRE1α-dependent unconventional splicing, leading to an increase of mScarlet/EGFP ratio ^49, 50^. When driven by the pan-motor neuronal driver, Tg[mnr2b-hs:Gal4], the mScarlet/EGFP ratio was higher in the large somas (Fig. 3g, h), suggesting that larger SMNs experienced higher levels of ER stress than small SMNs under normal conditions. To determine whether the UPR activation is specific to the IRE1α-Xbp1 pathway, we examined the status of the PERK-ATF4 pathway in the SMNs. Based on the principle of the SPOTlight reporter ^51^, we established a UAS reporter line (Tg[UAS:zSPOTlight]) to monitor the PERK-ATF4 pathway activity in zebrafish. Briefly, Gal4-dependent transcripts encoding the green (EGFP) and red (mScarlet) fluorophores were translated in a mutually exclusive manner depending on the activation of the PERK-atf4a pathway (Fig. 3i). This regulation is mediated by the differential usage of upstream open reading frames (uORFs) in the 5’ untranslated region of Atf4 mRNA ^52^, with mScarlet translation being enhanced under ER stress. When driven by the pan-motor neuronal driver, Tg[mnr2b-hs:Gal4], the mScarlet/EGFP ratio was significantly increased in the large SMNs (Fig. 3j, k). This indicates that the PERK-atf4a pathway was significantly active in large SMNs. Taken together, these results suggest that large SMNs inherently accumulate high levels of misfolded proteins in the ER, which may contribute to the global and inherent activation of cellular degradation systems.

### Accelerated cellular degradation is neuroprotective in large SMNs

To investigate whether enhanced cellular degradation is beneficial or detrimental to SMNs, we examined the consequences of autophagy perturbation in SMNs. To specifically manipulate autophagy in CaP, we targeted the expression of an ATG5 mutant with the lysine-to-arginine mutation (K132R), which disrupts the conjugation of ATG12 to ATG5, a critical step in autophagosome formation ^53^. ATG5^K132R^ and GFP-LC3-RFP-LC3ΔG were simultaneously expressed by bi-directional Gal4/UAS-mediated expression (Fig. 4a). ATG5^K132R^ overexpression significantly increased the GFP-LC3/RFP-LC3ΔG ratio not only in the CaPs but also in RB cells and interneurons at 48 hpf (Fig. 4b, c), confirming an inhibitory effect of ATG5^K132R^ on autophagic flux. Overexpression of wild-type ATG5 also inhibited autophagic flux in CaPs and RB cells, albeit to a lesser extent. Accordingly, we found that the total length of motor axons expressing wild-type ATG5 was reduced by an average of 7% and that the inhibitory effect was further exaggerated by the K132R mutation, resulting in a 24% reduction (Fig. 4d, e). Furthermore, CaP axons overexpressing ATG5^K132R^ were less ramified, although they still innervated their inherent target muscles in the ventral myotome. Based on these observations, we concluded that enhanced cellular degradation promotes motor axon outgrowth and is thus neuroprotective in large SMNs.

## Discussion

Through single-cell resolution scanning of autophagy, UPS, and UPR activities in the zebrafish spinal cord, we found that large SMNs inherently experience high catabolic stress and sustain the highest levels of cellular degradation activity in the spinal cord. These findings suggest that the junctions between the CNS and peripheral muscles are under significant catabolic stress even in healthy animals. This intrinsic vulnerability of the CNS-muscle connection may contribute to susceptibility to ALS in humans.

The intrinsically accelerated cellular degradation of large SMNs is a neuroprotective response to catabolic stress, as the disruption of ATG5-dependent autophagy leads to deficits in motor axon outgrowth. This finding aligns with a previous observation that motor neuron-specific ATG7 deletion in mice results in some denervated endplates in the tibialis anterior muscle, which are primarily innervated by fast-fatigable motor neurons ^21^. One potential reason autophagy positively influences motor axon outgrowth is its role in active metabolic turnover. Large SMNs, including fast-fatigable motor neurons, exhibit an active cellular metabolism to sustain their connections with muscles, regulate contractions, and provide trophic support ^38, 54^. These motor circuit functions are underpinned by the formation, maintenance, and elimination of synapses^21, 24, 27, 42, 55^, which are high energy-demanding subcellular compartments ^56^. Thus, the impairment of autophagy-dependent metabolic turnover could disrupt synaptic development and function, ultimately halting axonal growth in motor neurons. Beyond the role of this process in metabolic control, our findings are also open to the possibility that autophagy has a specialized function in degrading unidentified key proteins that inhibit motor axon outgrowth. The further amplification of the already elevated autophagy and UPS activity following the loss of TDP-43 function in large SMNs supports the idea that degradation substrates accumulate abnormally during ALS progression, compelling SMNs to upregulate their degradation machinery. TDP-43 is an RNA-binding protein whose primary functions include RNA splicing. Recent studies have demonstrated that loss of TDP-43 function leads to the incorporation of cryptic exons into mature mRNA ^43, 57–60^, resulting in the production of proteins with low folding fidelity ^48^. UPR imaging further indicated that misfolded protein accumulation is already evident under normal physiological conditions in large SMNs. Elevated ER stress has also been previously described in an SOD1-ALS model (SOD1-G93A mice) ^61, 62^ and in patients with sporadic ALS ^61^. Moreover, not only do misfolded proteins accumulate but also damaged organelles including mitochondria, owing to reduced TDP-43 function ^63^. Taken together, the increased degradation load from accumulated misfolded proteins and damaged organelles may be a common feature of both normal physiology and the pathophysiology of ALS-vulnerable neurons. In neurons that innately experience high degradation stress, risk factors affecting the cellular degradation systems, such as genetic mutations in autophagy and UPS-related components, may readily amplify proteostatic stress and reduce the threshold for degeneration ^64^. This type of vulnerability may share underlying mechanisms with the motor neurons that are lost during normal aging ^6^. Notably, the accumulation of autophagosomes in SMNs has been described in sporadic ALS cases and animal models of ALS ^18–20^, implying that the impairment of autophagy may be an inevitable outcome of TDP-43 loss, as the expression of some autophagy genes can be influenced by TDP-43 ^44, 57^. In such cases, the upregulation of autophagy and the UPS likely represents an early response of SMNs to a decline in TDP-43 function, which serves as the first line of defense to maintain proteostasis. This defense mechanism may eventually become overwhelmed by the increasing catabolic load over time, as autophagy gradually declines owing to the loss of TDP-43 function. We observed a decline in the UPS activity in a small population of CaP cells in TDP-43 DKO fish, which may reflect the sequential bidirectional effect of TDP-43 loss. In these scenarios, mitigating catabolic stress may be more fundamental and effective in preventing disease initiation than merely enhancing the degradation machinery.

A key question arising from the present study is why the catabolic load accumulates more in large SMNs than in small SMNs. Given that both the autophagy and the UPS are accelerated in large SMNs and can be further activated by the loss of TDP-43, dysfunction of the degradation system is unlikely to be the primary driver of degradation substrate accumulation. Instead, an increase in production may be a plausible explanation. SMNs are among the largest cell types in the body, allowing them to connect distant tissues, namely, the CNS and peripheral muscles, and transmit information over long distances. As a general characteristic of large-sized cells ^65^, high protein synthesis rates are assumed to increase the likelihood of misfolding, while oxidative stress from active metabolism disrupts protein folding, and a large cell volume dilutes protein quality control factors ^66, 67^. Clarifying how these vulnerabilities intrinsic to large cells are compensated for large SMNs is crucial for understanding the proteostatic control of both innate and disease-induced catabolic loads. The mechanism also provides insights into the design of therapeutic strategies to protect large SMNs from degeneration by alleviating the burden of cellular degradation.

Even though this study benefits from the ability to monitor proteostasis dynamics with cellular resolution in real time within the intact CNS, the approach is limited to zebrafish at the larval stage, when their tissues are highly translucent, allowing non-invasive and wide-field optical access to the CNS. Given that aging is a risk factor for ALS, further technological advances are needed to analyze the dynamics of autophagic flux, UPS, and UPR across the adult spinal cord segment to further validate the findings of this study. Nonetheless, this study provides important *in vivo* evidence of the proteostatic stress experienced by motor neurons under physiological conditions, which may drive the transition to pathological states.

## Methods

### Animals

This study was performed in accordance with the Guide for the Care and Use of Laboratory Animals of the Institutional Animal Care and Use Committee (IACUC) of National Institute of Genetics (NIG, approval numbers: 30-15, 31-14, R2-6, R3-1, R4-7, R5-1, R6-9, R6-10, R6-22). All experimental protocols were approved by the IACUC of NIG. All authors complied with the ARRIVE guidelines. Wild-type and transgenic zebrafish, hybrids of the AB and Tübingen strains, were used in all experiments. The fish were raised at 28°C under a 12-hour light/12-hour dark cycle (L/D) for the first five days post-fertilization. At these developmental stages, sex is not yet determined. Transgenic zebrafish lines used in this study were described in detail in Supplementary information.

### Mutant fish lines used in this study

For the generation of *tfeb* knockout fish, target sequences for Cas9-mediated cleavage were searched by CRISPRscan ^68^. The target sequences CGGTTTGAGCCTGGACCATGggg and TGGAGAGTGCATGTTCGGTGggg, where the protospacer adjacent motifs (PAMs) are indicated by lower cases, were chosen for the generation of *tfeb-n115* allele. A mixture of these two crRNAs and Streptococcus pyogenes Cas9 Nuclease V3 (Alt-R^®^ CRISPR-Cas9 System, IDT) was injected into one-cell stage zebrafish embryos. For the generation of *tfe3a* knockout fish, a mixture of two crDNAs (CGGAGCCGGTTGGGGTCACAggg and GGGCTTTGCGGGCAGGTTGGagg) was used. For the generation of *tfe3b* knockout fish, a crRNA (TAATCTAGGCTCCAACAAAGagg) was used.

### MG132 treatment

Zebrafish embryos at 30 hpf were treated with 100 µM MG132 or DMSO in E3 buffer at 28 °C for 24 hpf and analyzed by confocal microscopy at 54 hpf. The final concentration of DMSO was 1% in both treated and control conditions.

### Microscopic and statistical analyses

All images were acquired from live fish embedded in 0.8–1% low-melting agarose (NuSieve GTG Agarose, Lonza) on a Glass Base dish (IWAKI, 3010-035) using an Olympus FV1200 laser confocal microscope with a 20x water immersion objective (NA1.0). For confocal imaging, the fish were raised in an embryonic buffer containing 0.003% (w/v) N-phenylthiourea (SIGMA, P7629) to inhibit melanogenesis. Confocal images were acquired as serial sections along the z-axis, analyzed using Olympus Fluoview Ver2.1b Viewer and Image J, and processed for presentation using Adobe Photoshop and Illustrator. Statistical analyses were performed using GraphPad Prism Software.

## Acknowledgments

The authors are grateful to Drs Noboru Mizushima and Hideaki Morishita for sharing the GFP-LC3-RFP-LC3ΔG plasmid and instructing autophagic flux measurements and to Drs Haruhisa Inoue and Keiko Imamura for valuable discussions. This work was supported by The Nakabayashi Trust for ALS Research (K.A.), THE KATO MEMORIAL TRUST FOR NAMBYO RESEARCH (K.A.), Daiichi-Sankyo Foundation of Life Science (K.A.), Takeda Science Foundation (K.A.), KAKENHI Grant numbers JP22H04657 (K.A.), JP19K06933 (K.A.), JP22H02958 (K.A.), JP23H04266 (K.A.), JP21H02463 (K.K.), JP23H00375 (Y.S.) and AMED-PRIME under Grant Number JP23gm6410011h0003 (K.A.), and National BioResource Project (NBRP).

## Author contributions

K.A. conceived and designed the research. K.A. and S.S. performed the experiments. T.T and Y.S. provided the UPS probe. K.A., Y.S., TY and K.K. analyzed the data. K.A. and Y.S. wrote the manuscript with support from K.K. and H.H.. All authors discussed the results and commented on the manuscript.

## Extended Data

**Extended Data Fig. 1.**
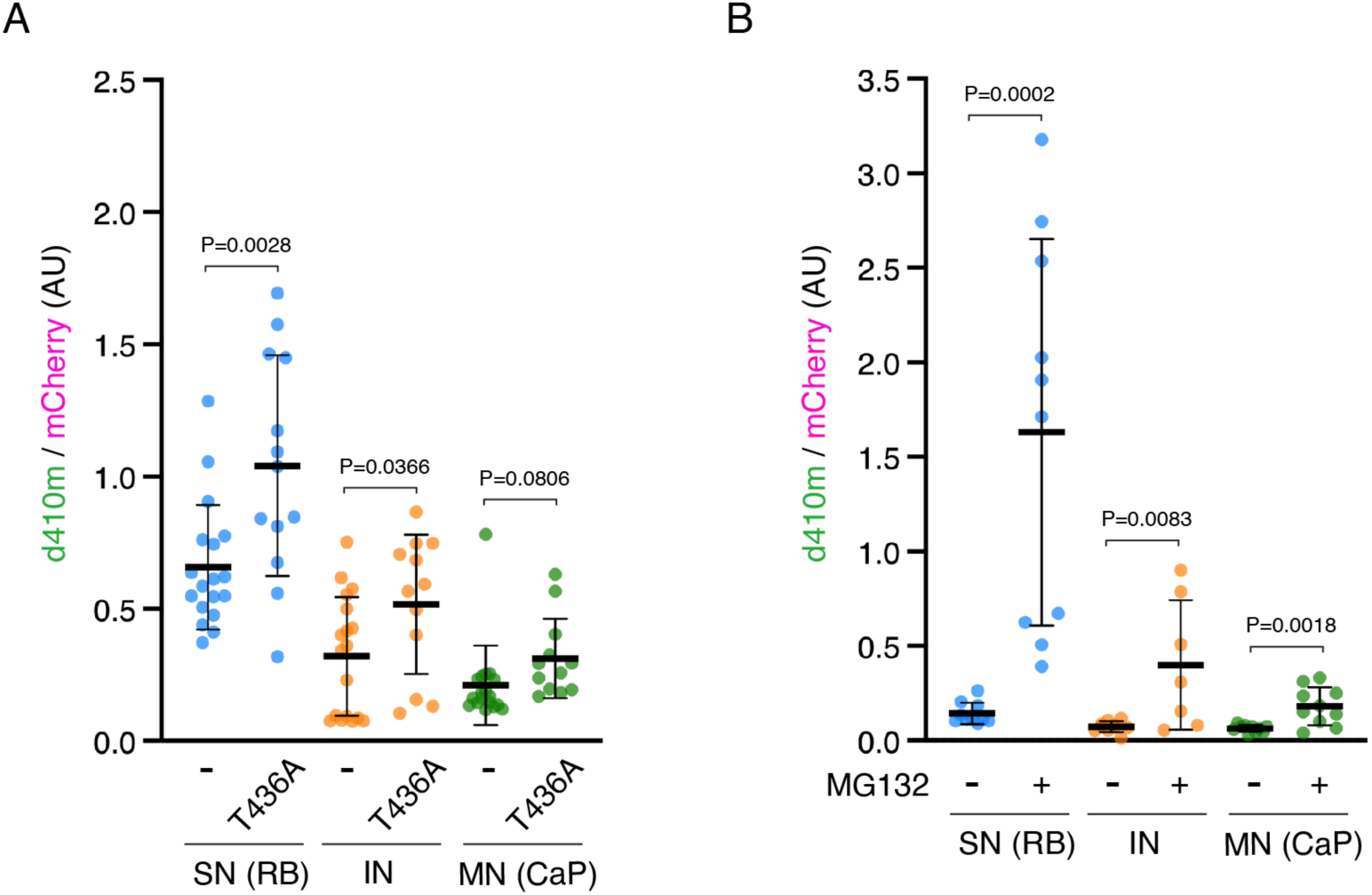
Development of the UPS reporter. (a) Comparison of stability between EGFP-d410m and EGFP-d410m-T436A. EGFP-d410m/mCherry (-) and EGFP-d410m-T436A/mCherry (T436) ratios in RB cells (SN), interneurons (IN) and CaPs (MN) were compared between Tg[SAIG213] Tg[UAS:hmCP2AGd410m] and Tg[SAIG213] Tg[UAS:hmCP2AGd410m-T436] fish at 30 hpf. Error bars show SD. (b) EGFP-d410m-T436A/mCherry (T436A) ratios in RB cells (SN), interneurons (IN) and CaPs (MN) were compared between Tg[SAIG213] Tg[UAS:hmCP2AGd410m] fish with (+) or without (-) MG132 treatment during 30 hpf - 54 hpf.

**Supplementary Video 1 Autophagic flux in the spinal cord**

Ratio images of spinal cord sections of Tg[SAGFF73A] Tg[UAs:GFP-LC3-RFP-LC3ΔG] shown in Fig. 1b (From medial to lateral). The bar indicates 20 µm.

