## Supplementary information for "Intrinsic and TDP-43 dysfunction-induced catabolic stress elicit neuroprotective cellular degradation in ALS-vulnerable motor neurons"

**RESOURCE AVAILABILITY**

Contact: Kazuhide Asakawa

Materials availability: Kazuhide Asakawa

Data availability: Kazuhide Asakawa

**Methods**

*Animals*

This study was performed in accordance with the Guide for the Care and Use of Laboratory Animals of the Institutional Animal Care and Use Committee (IACUC) of National Institute of Genetics (NIG, approval numbers: 30-15, 31-14, R2-6, R3-1, R4-7, R5-1, R6-9, R6-10, R6-22). All experimental protocols were approved by the IACUC of NIG. The all authors complied with the ARRIVE guidelines. Wild-type and transgenic zebrafish, hybrids of the AB and Tubingen strains, were used in all experiments. The fish were raised at 28°C under a 12:12 light/dark cycle (L/D) for the first five days post-fertilization. At these developmental stages, sex is not yet determined.

*The list of transgenic zebrafish lines used in this study*

Tg[UAS:GFP-LC3-RFP-LC3ΔG] (This study)

Tg[SAGFF73A] [1]

Tg[SAIG213A] [2]

Tg[mnr2b-hs:Gal4] [3]

Tg[UAS:tfebG-T2A-H2AmC] (This study)

Tg[UAS:tfe3aG-T2A-H2AmC] (This study)

Tg[UAS:tfe3bG-T2A-H2AmC] (This study)

Tg[ATG5:UAS:GFP-LC3-RFP-LC3ΔG] (This study)

Tg[ATG5K132R:UAS:GFP-LC3-RFP-LC3ΔG] (This study)

Tg[UAS:hmCP2AGd410m] (This study)

Tg[UAS:hmCP2AGd410mT436A] (This study)

Tg[UAS:Gxbp1uS] (This study)

Tg[UAS:zSPOTlight] (This study)

*The list of zebrafish mutant lines used in this study*

*tardbp-n115* [4]

*tardbpl-n94* [4]

*tfeb-n115* (This study)
*tfe3a-n141* (This study)
*tfe3b-n134* (This study)

*Generation of transgenic zebrafish lines*
All transgenic lines were generated by *Tol2* transposon-mediated transgenesis [5].

Tg[UAS:GFP-LC3-RFP-LC3ΔG]
The BamHI-BamHI fragment containing GFP-LC3-RFP-LC3ΔG was excised from the plasmid (a gift from the Mizushima Lab) [6] and introduced into the BglII site of pT2MUASMCS [2] to construct Tg[UAS:GFP-LC3-RFP-LC3ΔG].

Tg[ATG5:UAS:GFP-LC3-RFP-LC3ΔG], Tg[ATG5K132R:UAS:GFP-LC3-RFP-LC3ΔG]
The *Tol2* transposon cassette in which the amino (N)-terminally myc tagged zebrafish *atg5* (GenBank accession number NM_001009914) and GFP-LC3-RFP-LC3ΔG fragment were placed on both sides of the 5x upstream activation sequence (UAS) in an opposite direction was designed and synthesized (GeneArt Gene Synthesis, Thermo Fisher Scientific) to produce Tg[ATG5:UAS:GFP-LC3-RFP-LC3ΔG] line. The same construct except that the K132R (AAA to AGA) mutation was introduced into *atg5* was synthesized to generate Tg[ATG5K132R:UAS:GFP-LC3-RFP-LC3ΔG] line.

Tg[UAS:tfebG-T2A-H2AmC], Tg[UAS:tfe3aG-T2A-H2AmC], Tg[UAS:tfe3bG-T2A-H2AmC]
The *Tol2* transposon cassette carrying the 5x upstream activation sequence (UAS), the *hsp70l* promoter (650 bp), the open reading frame (ORF) of zebrafish *tfeb* (ENSDART00000006580_8) fused to EGFP at the carboxy (C)-terminus, a fragment encoding the T2A peptide [7], the ORF of *h2az2a* (GenBank accession number NM_153644) tagged with mCherry at the C-terminus, and the SV40 ployadenylation sequence, was synthesized (GeneArt Gene Synthesis, Thermo Fisher Scientific), to generate Tg[UAS:TFEBG-T2A-H2AmC]. Tg[UAS:tfe3aG-T2A-H2AmC] and Tg[UAS:tfe3bG-T2A-H2AmC] were constructed in the same manner as described above, except that the ORFs of *tfe3a* (GenBank accession number NM_131848) and *tfe3b* (GenBank

accession number NM_001045066) were used, respectively, instead of the *tfeb* ORF.

Tg[UAS:hmCP2AGd410m], Tg[UAS:hmCP2AGd410mT436A]
The mCherry-P2A-EGFPd410m fragment encoding mCherry linked via P2A peptide to EGFP tagged with the carboxy-terminus 52 amino acids of the mouse ornithine decarboxylase (GenBank accession number AAA51638) (EGFPd410m) was placed downstream the 5xUAS and *hsp70l* promoter (650 bp) and cloned into the *Tol2* transposon cassette to generate Tg[UAS:hmCP2AGd410m] line. The genetic mutation replacing Thr436 with Ala (ACG > GCG) in the ornithine decarboxylase is introduced by PCR-based mutagenesis to generate Tg[UAS:hmCP2AGd410mT436A] line.

Tg[UAS:Gxbp1uS]
The *Tol2* transposon cassette carrying the 5xUAS, the upstream and 5' untranslated regions (UTRs) of *gapdh* (GenBank accession number NM_001115114) (500 bp), the EGFP-mScarlet fusion linked with the 239-bp fragment of *xbp1u* (GenBank accession number BC044133) including 26 bp-stress dependent intron [8], and the SV40 ployadenylation sequence was synthesized (GeneArt Gene Synthesis, Thermo Fisher Scientific), to generate Tg[UAS:Gxbp1uS].

Tg[UAS:zSPOTlight]
The *Tol2* transposon cassette carrying the 5xUAS, uORF1 and uORF2 of *atf4a* (GenBank accession number NM_213233), EGFP, P2A, myc-tagged mScarlet and the SV40 ployadenylation sequence was synthesized (GeneArt Gene Synthesis, Thermo Fisher Scientific), to generate Tg[UAS:zSPOTlight]. In Tg[UAS:zSPOTlight], EGFP was inserted after the initiation codon of uORF2. mScarlet was inserted after the initiation codon of *atf4a*. The structure of this reporter is based on the SPOTlight reporter [9].

*Generation of zebrafish mutant lines*
For the generation of *tfeb* knockout fish, target sequences for Cas9-mediated cleavage were searched by CRISPRscan [10]. The target sequences CGGTTTGAGCCTGGACCATGggg and TGGAGAGTGCATGTTCGGTGggg, where the protospacer adjacent motifs (PAMs) are indicated by lower cases,

were used for the generation of tfeb-n115 allele. A mixture of these two crDNAs and Streptococcus pyogenes Cas9 Nuclease V3 (Alt-R® CRISPR-Cas9 System, IDT) was injected into one-cell stage zebrafish embryos. For the generation of *tfe3a* knockout fish, a mixture of two crDNAs (CGGAGCCGGTTGGGGTCACAggg and GGGCTTTGCGGGCAGGTTGGagg) was used. For the generation of *tfe3b* knockout fish, a crRNA (TAATCTAGGCTCCAACAAAGagg) was used.

The primers used for detection of the mutation by PCR followed by heteroduplex mobility assay (HMA) were
tfeb_checkF1: 5'-AAGGAGTACCTGTCCACCACATA-3'
tfeb_checkR1: 5'-CACTGTTTCCAGCTGAGTTCAT-3'
tfe3a_checkF1: 5'-AAGCTGATTTACCTTCTCATGGTC-3'
tfe3a_checkR1: 5'-ACTGCAGATTTAGGATTTGGTCAT-3'
tfe3b_checkF1: 5'-GGTGTAGGTGCAAACCCATC-3'
tfe3b_checkR2: 5'-CATAGGATGGCTGTGCTATGAA-3'
tardbp_HMAsg3sg4_F3: 5'-GCCAGATAATAAGAGGAAGATGGA-3'
tardbp_HMAsg3sg4_R3: 5'-TGACAGTACAAAGACAAACACCAC-3'
tardbpl_HMAsg3sg4_F2: 5'-CAATCACTGAATGAATGCACTTTT-3'
tardbpl_HMAsg3sg4_R2: 5'-GTTTGCTTATACTAACCTGCACCA-3'

*MG132 treatment*

Zebrafish embryos at 30 hpf were treated with 100 μM MG132 or DMSO in E3 buffer at 28 ˚C for 24 hpf and analyzed by confocal microscopy at 54 hpf. The final concentration of DMSO was 1% in both treated and control conditions.

*Microscopic and statistical analyses*

All images were acquired from live fish embedded in 0.8–1% low-melting agarose (NuSieve GTG Agarose, Lonza) on a Glass Base dish (IWAKI, 3010-035) using an Olympus FV1200 laser confocal microscope with a ×20 water immersion objective (NA1.0). For confocal imaging, the fish were raised in an embryonic buffer containing 0.003% (w/v) N-phenylthiourea (SIGMA, P7629) to inhibit melanogenesis. Confocal images were acquired as serial sections along the z-axis, and analyzed using Olympus Fluoview Ver2.1b Viewer and Image J, and processed for presentation using Adobe Photoshop and Illustrator. Statistical analyses were performed using GraphPad Prism Software.

1 Asakawa, K. *et al.* Genetic dissection of neural circuits by Tol2 transposon-mediated Gal4 gene and enhancer trapping in zebrafish. *Proc Natl Acad Sci U S A* **105**, 1255-1260 (2008). https://doi.org:10.1073/pnas.0704963105
2 Asakawa, K. & Kawakami, K. The Tol2-mediated Gal4-UAS method for gene and enhancer trapping in zebrafish. *Methods* **49**, 275-281 (2009). https://doi.org:10.1016/j.ymeth.2009.01.004
3 Asakawa, K. & Kawakami, K. Protocadherin-Mediated Cell Repulsion Controls the Central Topography and Efferent Projections of the Abducens Nucleus. *Cell Rep* **24**, 1562-1572 (2018). https://doi.org:10.1016/j.celrep.2018.07.024
4 Asakawa, K., Handa, H. & Kawakami, K. Optogenetic modulation of TDP-43 oligomerization accelerates ALS-related pathologies in the spinal motor neurons. *Nat Commun* **11**, 1004 (2020). https://doi.org:10.1038/s41467-020-14815-x
5 Suster, M. L., Kikuta, H., Urasaki, A., Asakawa, K. & Kawakami, K. Transgenesis in zebrafish with the tol2 transposon system. *Methods Mol Biol* **561**, 41-63 (2009). https://doi.org:10.1007/978-1-60327-019-9_3
6 Kaizuka, T. *et al.* An Autophagic Flux Probe that Releases an Internal Control. *Mol Cell* **64**, 835-849 (2016). https://doi.org:10.1016/j.molcel.2016.09.037
7 Kim, J. H. *et al.* High cleavage efficiency of a 2A peptide derived from porcine teschovirus-1 in human cell lines, zebrafish and mice. *PLoS One* **6**, e18556 (2011). https://doi.org:10.1371/journal.pone.0018556
8 Li, J. *et al.* A transgenic zebrafish model for monitoring xbp1 splicing and endoplasmic reticulum stress in vivo. *Mech Dev* **137**, 33-44 (2015). https://doi.org:10.1016/j.mod.2015.04.001
9 Helseth, A. R. *et al.* Cholinergic neurons constitutively engage the ISR for dopamine modulation and skill learning in mice. *Science* **372** (2021). https://doi.org:10.1126/science.abe1931
10 Moreno-Mateos, M. A. *et al.* CRISPRscan: designing highly efficient sgRNAs for CRISPR-Cas9 targeting in vivo. *Nat Methods* **12**, 982-988 (2015). https://doi.org:10.1038/nmeth.3543
